# Longitudinal Profiling of CD4^+^ T Cell Responses Following de novo Yellow Fever Vaccination

**DOI:** 10.64898/2026.02.26.708120

**Authors:** Dzenis Gojak, Maria Kuznetsova, Vincent Van Deuren, Sami Alcedo, Elizabeth Willems, Hajar Besbassi, Ruben Bond, Esther Bartholomeus, Joachim Marien, Kevin Arien, Pieter Meysman, Patrick Soentjens, My Ha, Benson Ogunjimi

## Abstract

The yellow fever 17D vaccine is one of the most successful live-attenuated viral vaccines, yet the cellular mechanisms underlying its long-term protection remain not fully understood. This study provides a longitudinal analysis of the human CD4^+^ T cell and IgG response following de novo yellow fever vaccination by focusing.

Peripheral blood mononuclear cells (PBMCs) from 49 vaccinated individuals were stimulated with yellow fever (YF) and control peptide pools across four timepoints: pre-vaccination/baseline (D1), day 22 (D22), day 43 (D43), and one year (D365) post-vaccination. Activation-induced marker (AIM) assays confirmed robust activation of CD4^+^ T cells following yellow fever peptide stimulation, peaking at day 22 post-vaccination and subsequently declining. T-cell receptor (TCRβ) sequencing of AIM-sorted CD4^+^ T cells showed a transient increase in clonal diversity at D22, consistent with broad epitope targeting and early polyclonal expansion. This was followed by repertoire contraction, which could indicate the persistence of a limited set of dominant clonotypes responsible for immune memory formation. TCR repertoires remained largely private over time, indicating a mostly individualized immune response. Next, serological analyses revealed a robust and highly yellow fever virus (YFV)-specific IgG response. Antibody levels peaked within the D22–D43 window and remained elevated at one year post-vaccination. Cross-reactivity toward other flaviviruses was limited, suggesting an antigen-specific humoral response.

Together, these findings characterize the longitudinal dynamics of the CD4^+^ T cell and IgG response following de novo yellow fever vaccination and provide insights into the mechanisms contributing to durable antiviral immunity.

## Introduction

Live-attenuated vaccines provide a controlled and minimally invasive method to study human T cell dynamics in vivo. Vaccines have become a widely used model to investigate the kinetics, diversity, and long-term persistence of vaccine-induced T cell responses in humans [1,2,7]

The yellow fever (YF) 17D vaccine has been in use for decades and has greatly reduced the YF burden. However, vaccine-induced protection differs between individuals. This variation is influenced by host-specific factors such as genetics, age, immune history, and overall health status [4]. Until recently, most studies focused on antibodies as the main correlate of protection for assessing vaccine efficacy. However, studies into T-cell response to many of the modern vaccines has remained largely underrepresented. A more detailed understanding of how human T cells respond to viral vaccines over time could help establish key benchmarks for protective immunity and guide future vaccine design efforts.

## Results

### Peak CD4^+^ T Cell Activation at Day 22

49 healthy adults received de novo YF vaccination and were sampled at baseline, day 22, day 43 and day 365. To quantify vaccine-induced activation of CD4^+^ T cells, we performed an activation-induced marker (AIM) assay after stimulation with a collection of YF-peptides [5] and parvovirus B19 NS1 (parv) peptides as a control. AIM^+^ responses were defined as CD154^+^CD134^+^ for CD4^+^ T cells. All values were background-corrected for the unstimulated condition (neg).

The response to YFV stimulation peaked sharply at D22, with a mean AIM^+^ frequency exceeding 1% of the total CD4^+^ compartment (Figure 1). This response was significantly higher than both the D1 baseline and the parvovirus B19 control at the same timepoint (p < 0.0001). While AIM^+^ CD4^+^ frequencies declined at D43 and returned near baseline levels by D365, the overall pattern could be indicative of a classical activation-contraction dynamic consistent with effector expansion followed by memory formation. The parvovirus-specific activation remained low and stable over time, contrasting with the dynamic vaccine-induced YFV response.

**Figure 1.**
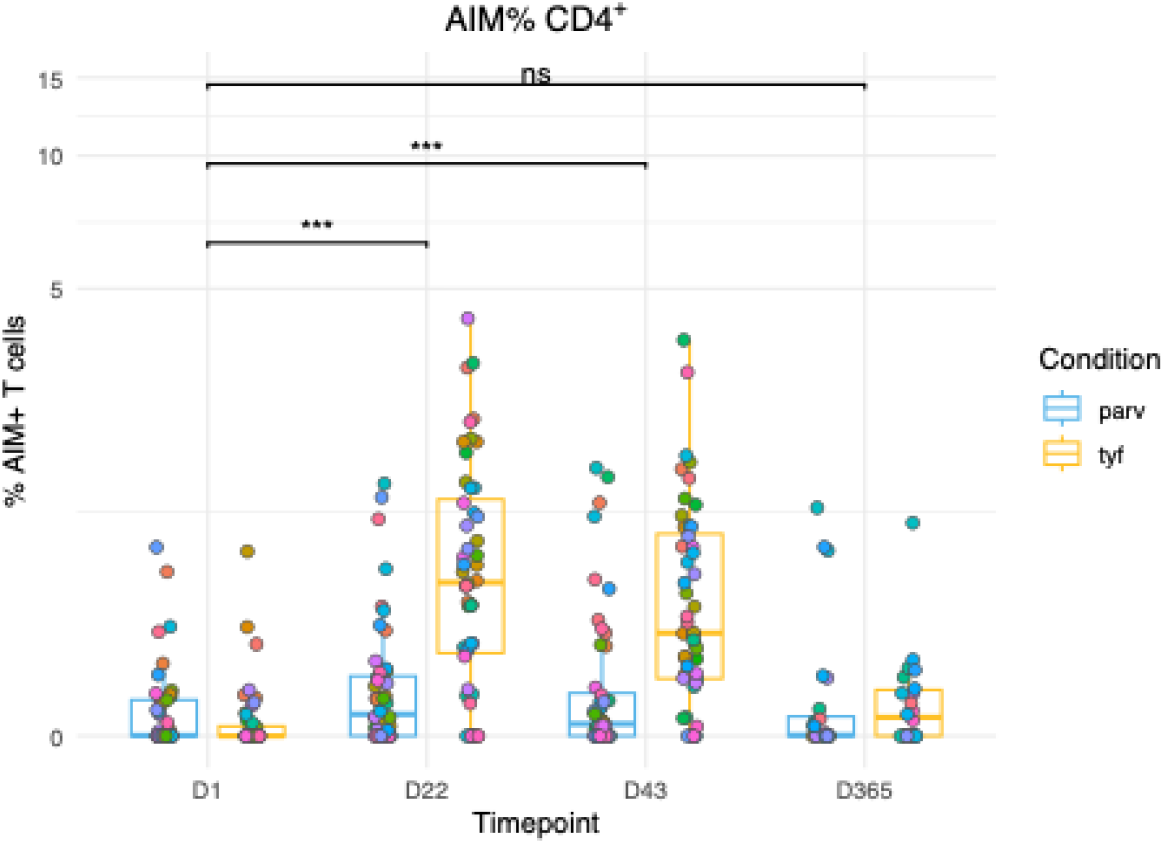
Antigen-specific activation of CD4^+^ T cells. Background-corrected Δ AIM% for CD4^+^ T cells following stimulation with yellow fever (tyf) or parvovirus B19 (parv) peptide pools at baseline (D1), Day 22 (D22), day 43 (D43), day 365 (D365) (n = 49, n = 38 for D365). Statistical significance between YFV-stimulated samples was determined using paired Wilcoxon signed-rank tests (*p ≤ 0.01, **p ≤ 0.001, ***p ≤ 0.0001, ns = not significant).

### Polyclonal Expansion Followed by Repertoire Contraction

Shannon entropy is widely used in TCR repertoire analysis as a measure of clonal diversity, capturing both the richness (number of unique clonotypes) and evenness (distribution of their frequencies) within a sample. It provides a single-value summary of repertoire complexity, allowing comparison across timepoints, cell types, or stimulation conditions. Higher entropy often indicates a more diverse repertoire, with many different clonotypes present, whereas lower entropy often indicates a less diverse repertoire of expanded clones, as typically seen in antigen-driven responses [6]. Because it accounts for both dominant and rare clones, Shannon entropy is particularly suitable for quantifying immune repertoire dynamics.

Absolute Shannon entropy values revealed a distinct difference between the AIM^+^ CD4^+^ T cell population and the negative control (unstimulated, D1 only) control group (Figure 2). Across all timepoints, the negative background consistently showed higher entropy values, suggesting a broader, unstimulated TCR repertoire, possibly caused by less antigen-driven specialization and the absence of dominant, epitope-specific clonotypes. In contrast, the AIM^+^ CD4^+^ subset showed lower entropy, consistent with clonal selection following antigenic stimulation. Notably, the overall pattern closely mirrors the AIM data, with a peak at D22 followed by contraction.

**Figure 2.**
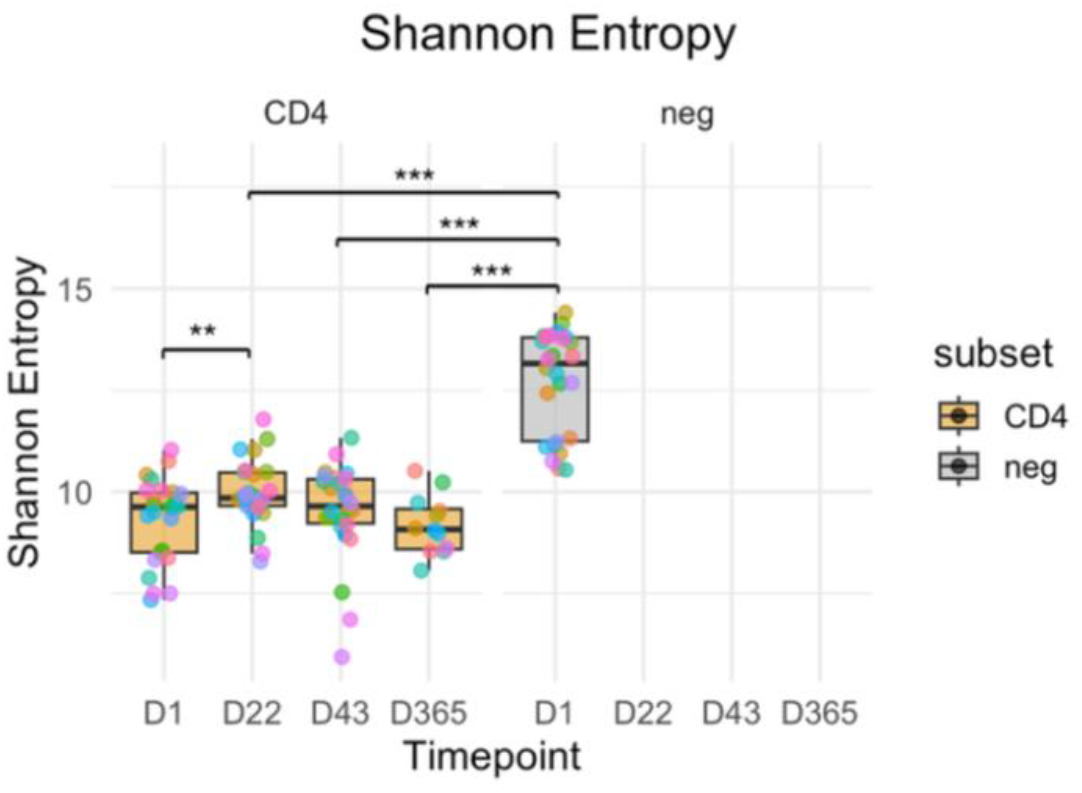
Absolute Shannon entropy across timepoints for AIM^+^ CD4^+^ T cell repertoires compared to the AIM^+^ (negative control) background (n = 26). Shannon entropy was calculated at baseline (D1), early (D22), intermediate (D43), and late (D365) post-vaccination. The AIM^−^ (neg) control group exhibited consistently higher entropy, reflecting a broader and unspecialized TCR repertoire. In contrast, the AIM^+^ CD4^+^ subsets showed reduced entropy consistent with antigen-driven clonal focusing. Colored dots represent individual participants. Statistical comparisons were performed using a paired Wilcoxon signed-rank test, with Benjamini– Hochberg correction for multiple comparisons (*p ≤ 0.01, **p ≤ 0.001, ***p ≤ 0.0001).

A paired Wilcoxon signed-rank test with Benjamini-Hochberg correction was used to assess if Shannon entropy differed significantly between the AIM^+^ group and the unstimulated control. The AIM^+^ CD4^+^ group showed significantly lower entropy compared to the negative control (p = 2.98 × 10^−8^ for both, Benjamini-Hochberg adjusted p = 1.19 × 10^−7^). These results suggest that YFV peptide stimulation narrows the T cell repertoire by selectively expanding a focused group of clonotypes, in contrast to the broader and more unspecialized background in the unstimulated control. These patterns are consistent with a broader, polyclonal expansion phase early after vaccination (D22), driven by the activation of many clones. The subsequent reduction in entropy suggests a narrowing of the repertoire, as high-affinity or immunodominant clones become selectively maintained or transition into memory pools.

To assess the degree of clonal persistence and repertoire stability over time, pairwise AIM^+^ TCR clone overlap was calculated between sequential timepoints using the percentage of shared clonotypes between donor-matched samples. Analyses were conducted for D1 vs D22, D22 vs D43, and D43 vs D365 using matrix visualizations. Clonal overlap between D1 and D22 was generally low, with most donors sharing less than 10% of clones across the timepoints (Figure 3). This observation may reflect turnover in the detectable TCR repertoire following vaccination, although it should be interpreted with caution given the inherently stochastic nature of TCR repertoire sampling and sequencing depth limitations, in addition to the emergence of vaccine-induced clonotypes that were absent or below detection thresholds at baseline. Between D22 and D43, the degree of overlap increased modestly in a subset of individuals, with several donors exhibiting 10–20% shared clones. This may indicate short-term persistence of dominant clones recruited during the peak effector phase, although overall overlap remained relatively modest, supporting a dynamic reshaping of the repertoire during the contraction and early memory phases.

**Figure 3.**
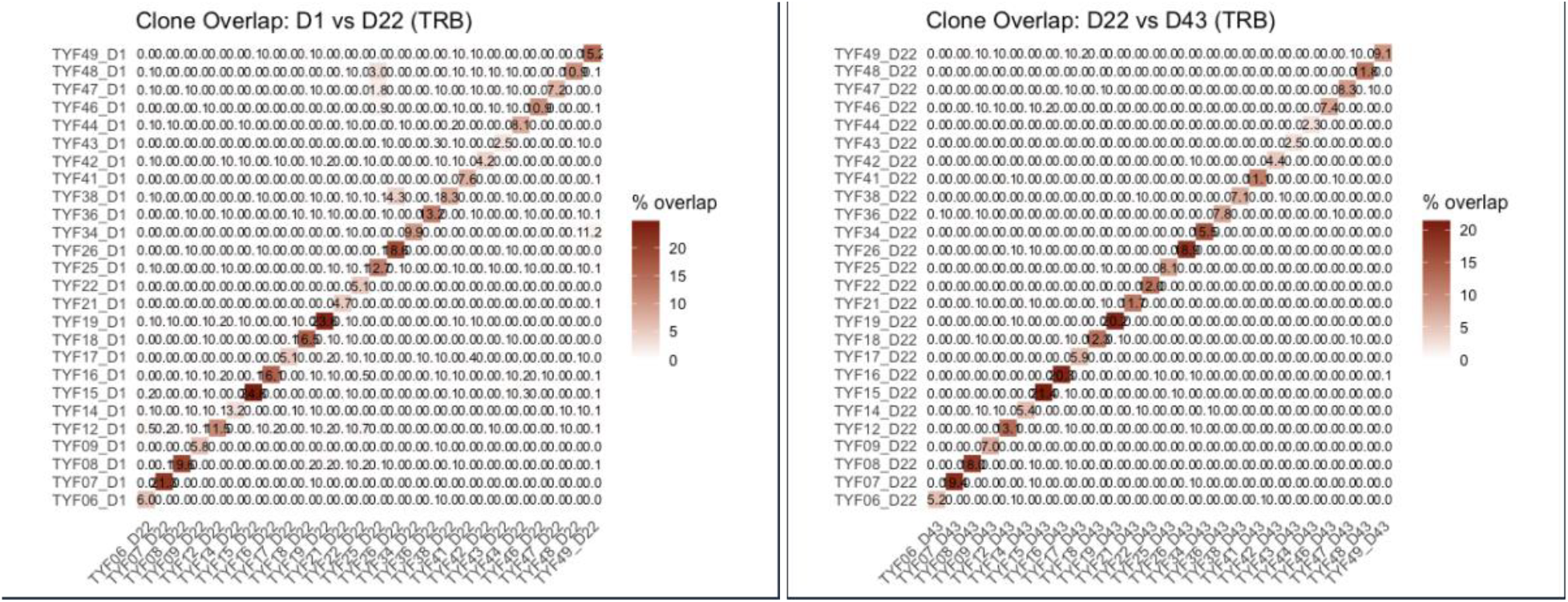
Clone overlap between adjacent timepoints: D1 vs D22 (left) and D22 vs D43 (right). Heatmaps show the percentage of shared TCR clones between donor-matched samples. Low overlap between D1 and D22 suggests de novo recruitment of vaccine-specific clonotypes, while higher overlap from D22 to D43 indicates short-term persistence of some expanded clones.

At the longest interval (D43 vs D365), a small number of participants retained a detectable fraction of clones (up to ∼16%) across timepoints, suggesting long-term persistence of certain vaccine-responding clonotypes (Figure 4). However, for most donors, overlap remained low or undetectable by one year post-vaccination. This pattern supports the conclusion that while some degree of clonal overlap persists, the overall TCR repertoire remains largely private.

**Figure 4.**
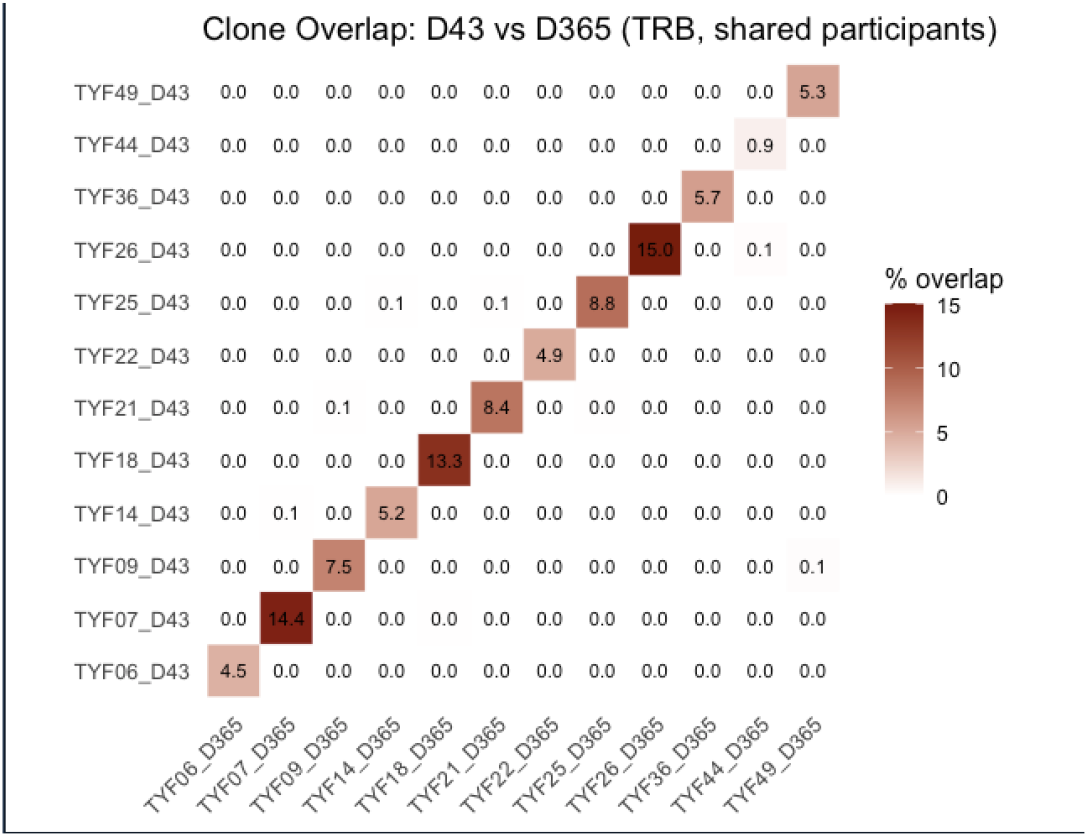
Clone overlap between D43 and D365 reveals long-term clonal persistence. Selected donors retained up to ∼16% of their D43 clones one year later, suggesting stable memory formation. However, most donors exhibited minimal or no overlap, highlighting the individualized nature of the TCR repertoire over time as a select group of dominant clones is responsible for long-term memory formation.

Across all timepoint comparisons, within-donor overlap consistently exceeded between-donor overlap, demonstrating the private nature of YFV vaccine-induced TCR repertoires. The symmetric heatmaps clearly illustrate this donor specificity, with overlap concentrated along the diagonal. Together, these findings indicate that YF vaccination induces a dynamic and individualized clonal response, with modest persistence of dominant clones over time. The low baseline-to-D22 overlap further supports the emergence of de novo vaccine-specific clonotypes, while the moderate D22-to-D43 and occasional D43-to-D365 retention hint at the formation of a selective memory pool.

These overlap patterns complement the entropy results by revealing how clonal turnover and memory formation shape the TCR repertoire post-vaccination. The transient rise in entropy at D22 aligns temporally with the peak in antigen-specific T cell activation observed in the flow cytometry data. This suggests a coordinated response characterized by the recruitment of diverse clonotypes during the early effector phase. The gradual reduction in entropy by D43 and D365 may reflect repertoire focusing during memory formation or selective persistence of a subset of vaccine-induced clones.

### Robust YFV-Specific IgG Responses Peak Between D22–D43

Longitudinal analysis of YFV-specific IgG responses (using Luminex xMAP multiplex bead-based immunoassay) revealed a consistent pattern of vaccine-induced induction across participants (Figure 5A, 5B, 5C). At baseline (D0), IgG levels were low and confined to a narrow range. Early timepoints (D2 and D8) showed minimal deviation from baseline. A clear upward shift can be observed by D15, followed by a further increase at D22 and D29. IgG levels remained elevated at D43 and were largely sustained at D365. However, noticeable interindividual variability in peak magnitude was observed. Some participants exhibited strong expansions exceeding 10^3^ fluorescence intensity (FI) units, whereas others showed more moderate increases. These results align temporally with both the AIM and TCR data, with a noticeable shift starting at D22 and continuing through D43.

**Figure 5A.**
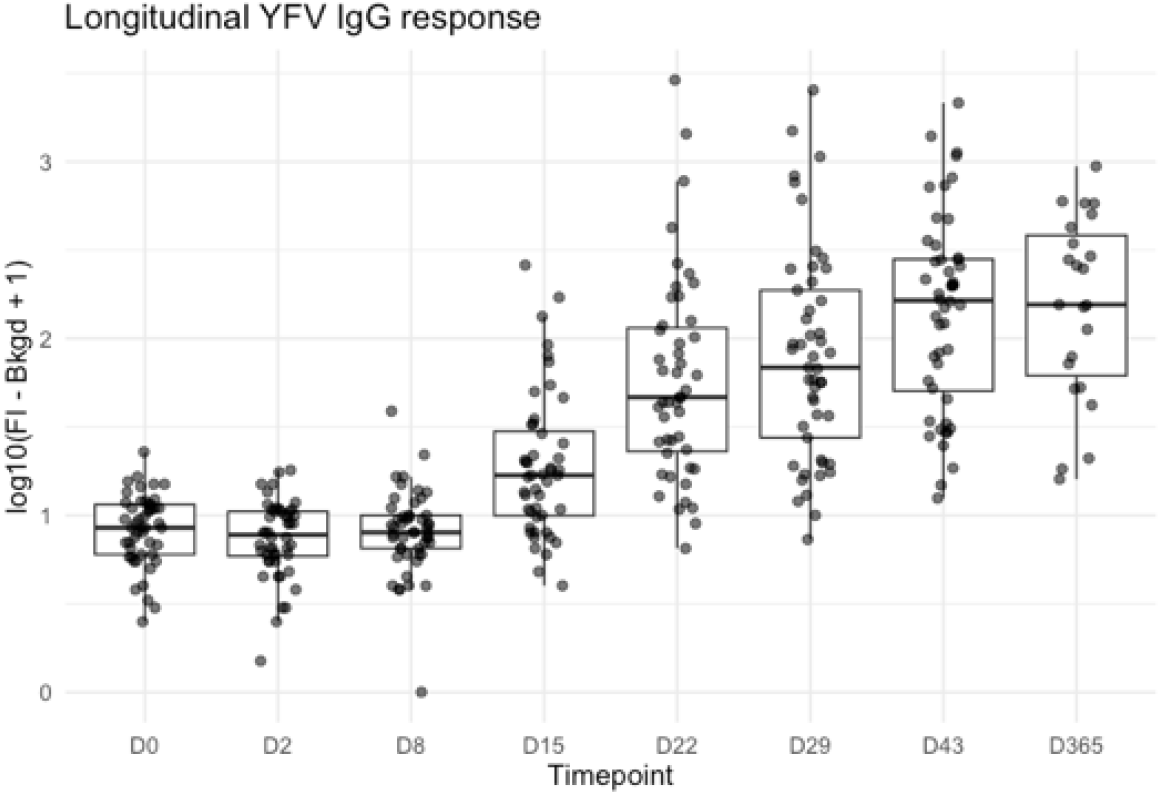
Longitudinal YFV IgG response (cohort-level). Boxplotsof log10(FI−Bkgd+1) YFV-specific IgG levels at each timepoint. Points represent individual participants. Median responses rise from D15 onward, peak at D22–D43, and remain elevated at D365.

**Figure 5B.**
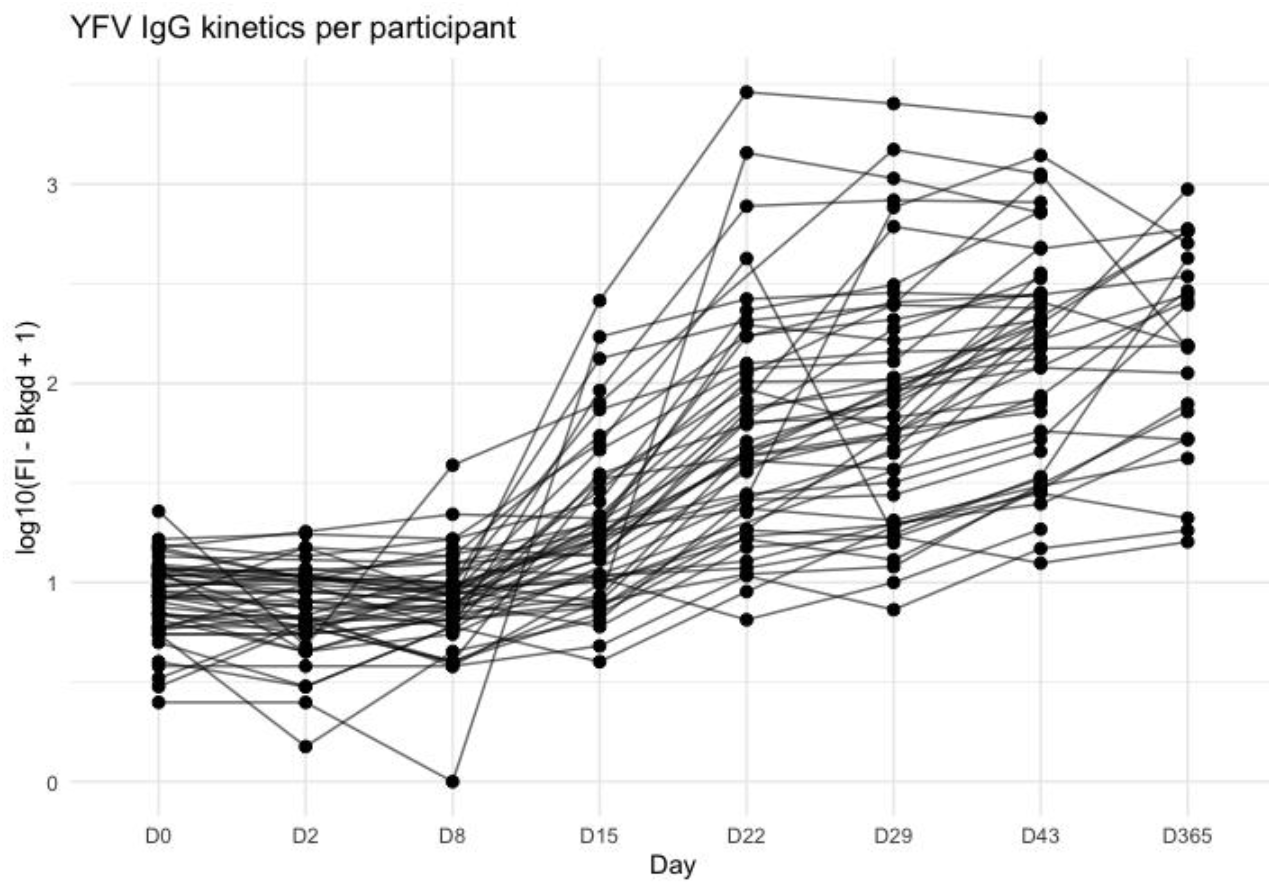
Longitudinal kinetics of YFV-specific IgG responses per participant. Individual trajectories of YFV-specific IgG levels across all timepoints (D0–D365). IgG values are shown as log10(FI−Bkgd+1). Each line represents one participant.

**Figure 5C.**
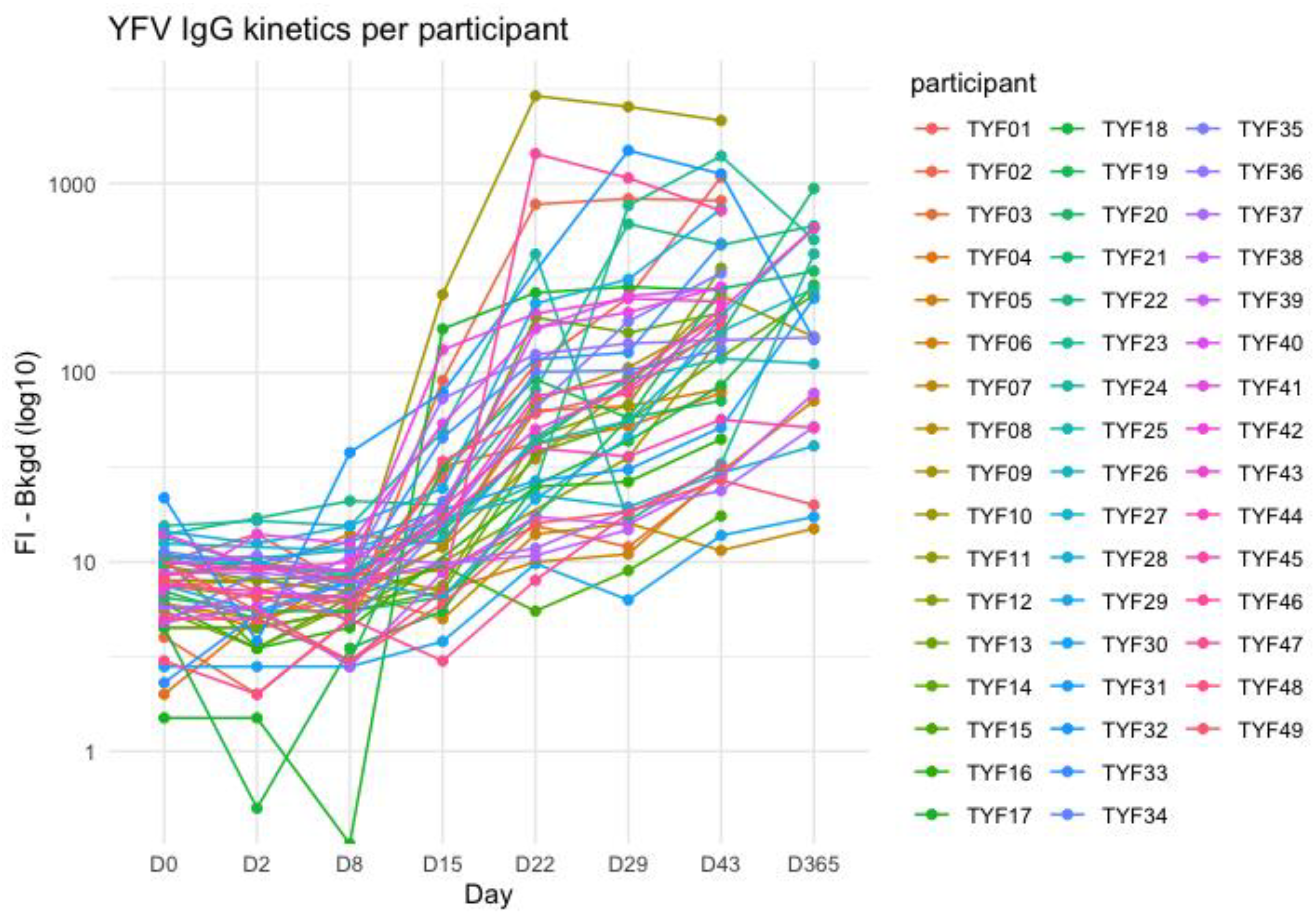
YFV IgG kinetics per participant. Individual trajectories of YFV-specific IgG (FI−Bkgd, log10 scale on y-axis) across all timepoints. Each line represents one participant. The figure illustrates delayed induction beginning around D15 and peak responses at D22–D43.

To determine whether the antibody response was YFV-specific rather than broadly flavivirus-reactive, antigen binding profiles were examined at each participant’s YFV peak day (Figure 6). For each participant, the YFV peak day was defined as the timepoint within the predefined peak window (D22–D43) at which FI for the YFV bead reached its maximum value. YFV-specific binding was significantly elevated compared to other flavivirus antigens. While modest increases in some related flaviviruses were detectable in certain individuals, these responses were substantially lower in magnitude. Figure 7 shows that YFV-specific induction was consistently strong across the cohort, whereas responses for other flaviviruses were generally modest or absent.

**Figure 6.**
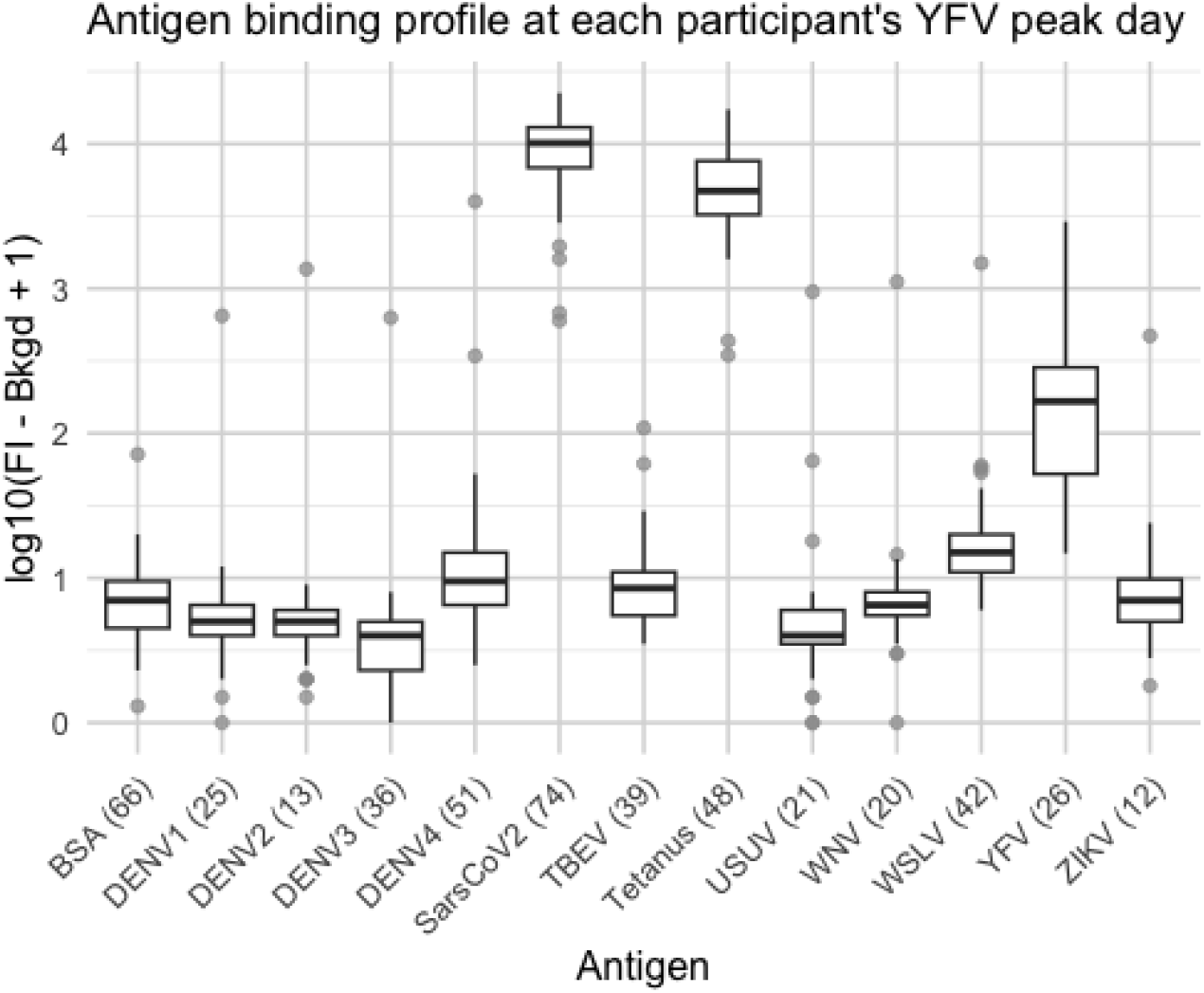
Antigen binding profile at each participant’s YFV peak day. Boxplots of log10(FI−Bkgd+1) binding to all panel antigens at the individual YFV peak timepoint. YFV-specific binding is noticeably higher than other flaviviruses.

**Figure 7.**
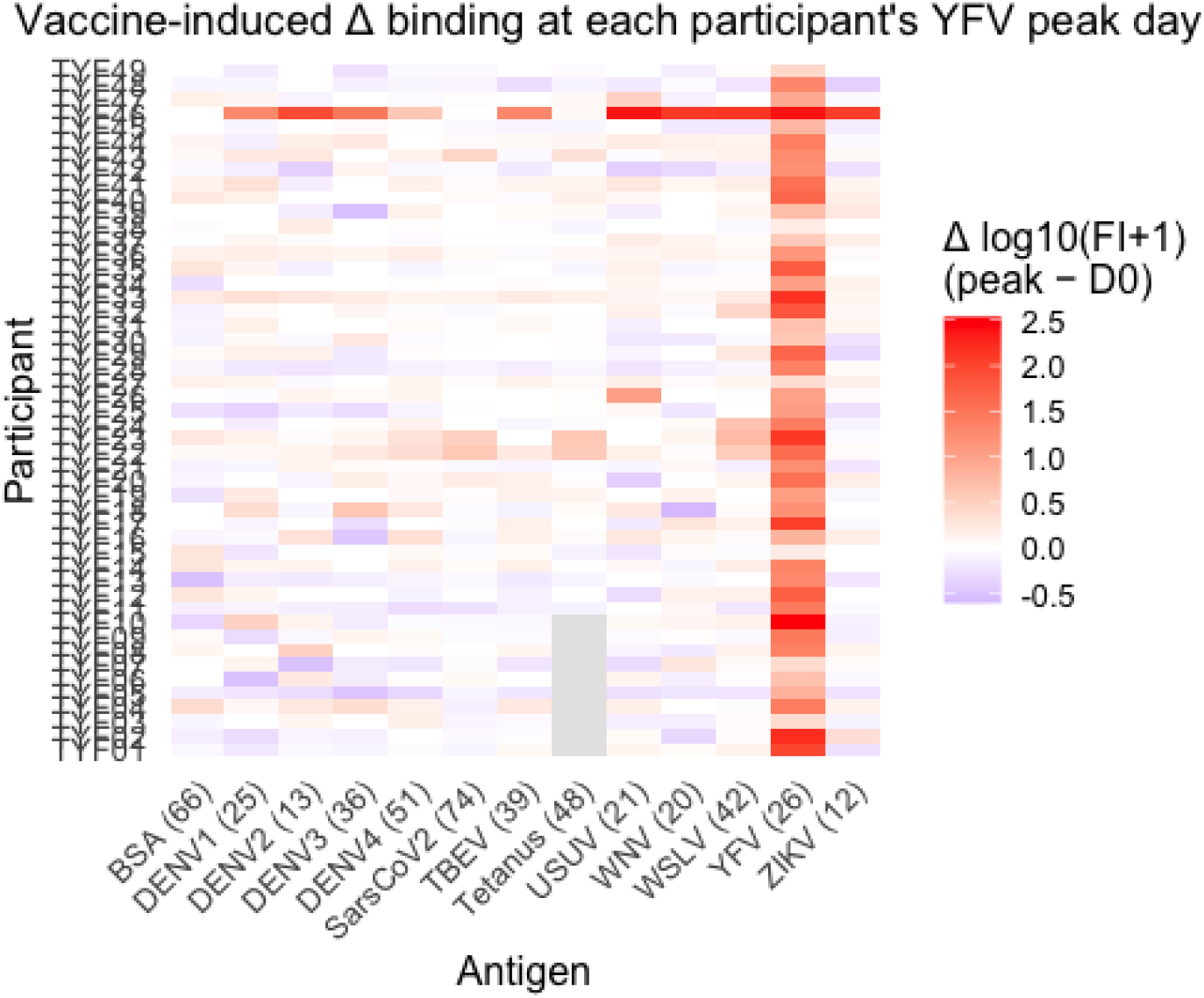
Vaccine-induced Δ binding at each participant’s YFV peak day. Heatmap of Δlog10(FI−Bkgd+1) (peak − D0) per antigen and participant. YFV-specific induction is consistently strong, whereas Δ responses for other flaviviruses are comparatively modest.

To analyze the IgG data, a linear mixed-effects model (LMM) was fitted with timepoint as a fixed effect and participant as a random intercept. The omnibus effect of timepoint was highly significant (p < 10^−70^), confirming that IgG levels varied substantially across the longitudinal study period. Baseline-referenced contrasts demonstrated that IgG levels did not differ significantly from D0 at D2 or D8 but were significantly elevated from D15 onwards (BH-adjusted).

For antigen specificity, paired non-parametric testing of delta responses confirmed that YFV-specific induction was significantly greater than the mean induction across other flaviviruses (p < 10^−8^), supporting the specificity pattern shown in figures 5 and 6.

## Discussion

The longitudinal analysis of AIM^+^ CD4^+^ T cells after de novo YF vaccination revealed that statistically significant changes were exclusively observed in the YFV-stimulated group. These results demonstrate that activation of CD4^+^ T cells occurs specifically in response to YFV peptide stimulation, particularly during the early post-vaccination window. The B19-stimulated group remained stable, lacking statistically robust changes in activation. These findings demonstrate the coordinated and time-sensitive nature of the YFV-induced immune response.

The observed peak in Shannon entropy at day 22 aligns with the timing of the strongest immune activation seen in the flow cytometry data, where CD4^+^ T cell activation markers were most prominently upregulated. This peak in entropy suggests an increase in clonal diversity within the TCRβ repertoire, which is typical for the early stages of a robust immune response. As shown in previous studies, including by Qi et al. [7], an initial rise in the number of epitope-specific T cells is a well-described feature following vaccination. Similarly, Pogorelyy et al. [9] observed a sharp increase in YFV-reactive clonotypes around day 15, suggesting a strong, early antigen-driven response. Although these studies measured increases in the abundance of epitope-specific clones, our data show a rise in Shannon entropy, possibly reflecting not just expansion but a broad and polyclonal recruitment of distinct T cell clonotypes.

This increase in diversity at day 22 may result from the simultaneous activation and expansion of many different epitope-specific clones. Over time, as a smaller number of high-affinity or better-adapted clones begin to dominate the response, overall repertoire diversity contracts at later timepoints (D43 and D365). Thus, the elevated Shannon entropy at day 22 likely captures a highly polyclonal and dynamic phase of the immune response just prior to repertoire narrowing and memory formation.

Regarding the IgG data, the results demonstrate a robust YFV-specific humoral response that is highly antigen-specific and displays only limited cross-reactivity toward other flaviviruses included in the panel. Additionally, the longitudinal profile closely resembles that of the cellular response, with minimal early changes, a clear induction phase beginning around D15, and peak responses between D22 and D43. This temporal pattern mirrors the expansion dynamics observed in the CD4^+^ AIM^+^ compartment and the clonal amplification detected in the TCR repertoire during the same window.

Together, these findings suggest coordinated kinetics between the humoral and cellular immune response following vaccination.

## Acknowledgments

We would like to thank all participants for their participation. We would like to thank Kato Van Bortel, Lida Van Petersen and Karin Peeters for their help in recruitment and sampling.

## Participants and Methods

### Participants

This longitudinal study included 49 healthy adult volunteers who received a single dose of de novo live-attenuated yellow fever 17D vaccine. Peripheral blood samples were collected at baseline prior to vaccination (D0) and different post-vaccination timepoints: day 2 (D2), day 8 (D8), day 15 (D15), day 22 (D22), day 43 (D43), and one year (D365). Only D0, D22, D43, and D365 were used in the AIM assay and TCR sequencing. Not all participants contributed samples at D365 due to follow-up availability. All participants provided written informed consent, and the study was conducted in accordance with institutional and national ethical guidelines (Antwerp University Hospital IRB number 15/19/210). Participant identifiers were pseudonymized prior to analysis.

### PBMC Processing and Cryopreservation

PBMCs were isolated from whole blood by density-gradient centrifugation using Lymphoprep (StemCell Technologies) in SepMate tubes (StemCell Technologies) according to the manufacturer’s instructions. Following isolation, cells were washed, counted, and assessed for viability. PBMCs were cryopreserved in fetal bovine serum supplemented with dimethyl sulfoxide using controlled-rate freezing and stored in liquid nitrogen until use. For downstream experiments, cryovials were rapidly thawed, washed to remove cryoprotectant, and rested in complete RPMI medium.

### Stimulation With Peptide Pools and Sorting

For antigen-specific activation assays, PBMCs were stimulated in vitro with overlapping peptide megapools spanning yellow fever virus (YFV) proteins comprising 15–20 amino acid peptides, with an average overlap of 9-10 amino acids, spread across structural proteins and non-structural proteins [5]. A parvovirus B19 peptide pool (GenScript Biotech) was included as an unrelated antigen control, and unstimulated wells served as negative controls. Cells were cultured overnight in the presence of co-stimulatory antibodies.

Following stimulation, cells were stained for viability and surface markers to identify activation-induced marker (AIM)–positive T cells. Antigen-specific CD4^+^ T cells were defined by co-expression of CD154 and CD134 after YFV peptide stimulation. For TCR repertoire analysis, AIM^+^ CD4^+^ T cells were isolated by fluorescence-activated cell sorting (FACS) using a Cytek Aurora CS spectral cell sorter (Cytek Biosciences). Baseline negative-control populations were also sorted to serve as background references. Sorted cells were collected in OMICS-Guard buffer and processed for downstream library preparation using the TIRTL-seq protocol as previously described [8].

### Flow Cytometric Analysis

Flow cytometric acquisition was performed to measure activation of CD4^+^ T cells. A standardized gating strategy was applied across all samples to identify AIM^+^ subsets. Antigen-specific responses were quantified as background-corrected ΔAIM%, calculated by subtracting the frequency observed in unstimulated wells from that in peptide-stimulated wells.

### Bulk TCR Sequencing

TCRβ repertoires were profiled from sorted AIM^+^ CD4^+^ T cells using a bulk template-switch–based library preparation approach incorporating unique molecular identifiers and dual sample indices. Library preparation was performed using the TIRTL-seq method [8], followed by high-throughput paired-end sequencing on an Element AVITI− system.

### Luminex xMAP multiplex bead-based immunoassay

IgG levels were quantified using a Luminex xMAP multiplex bead-based immunoassay, and all serological measurements are presented as log10(FI−Bkgd+1), where FI−Bkgd represents the fluorescence intensity of antigen binding after subtraction of background signal.

### Statistical Analysis

To ensure specificity, enrichment of AIM^+^ cells in YFV-stimulated versus unstimulated conditions was assessed using Fisher’s exact test for each donor and timepoint. Only statistically significant responses after false discovery rate (FDR) correction were retained for downstream analyses; non-significant responses were set to zero. Data from multiple acquisition runs were merged, quality controlled, and batch effects were evaluated and corrected computationally prior to statistical analysis.

Paired Wilcoxon signed-rank tests were used to compare delta AIM% between D1 and subsequent timepoints (D22, D43, D365) for each stimulus and to compare YF vs parvovirus B19 responses at each timepoint using matched donor samples. The Wilcoxon signed-rank test is a non-parametric alternative to the paired t-test and was chosen because it does not assume a normal distribution of the data.

Paired Wilcoxon signed-rank tests were used to compare Shannon entropy between D1 and subsequent timepoints (D22, D43, D365). All analyses were restricted to the TCRβ chain and based on clone definitions using the combination of V gene, nucleotide CDR3 sequence, and J gene.

P-values were corrected for multiple testing using the Benjamini-Hochberg method. In contrast to the AIM assay, the TCR repertoire comparisons involved a smaller number of predefined hypotheses, but the underlying metrics, such as Shannon entropy, were derived from complex and high-dimensional sequencing data. Due to the greater variability and biological noise associated with immune repertoire measurements, the Benjamini-Hochberg procedure was applied to correct for multiple testing. While it allows for a higher rate of false positives among significant findings, it ensures that the proportion of such false negatives remains limited. This approach is particularly appropriate for exploratory analyses of TCR diversity, where the aim is to identify meaningful changes across timepoints and subsets without overly reducing statistical power.

---

**Supplementary figure 1.**
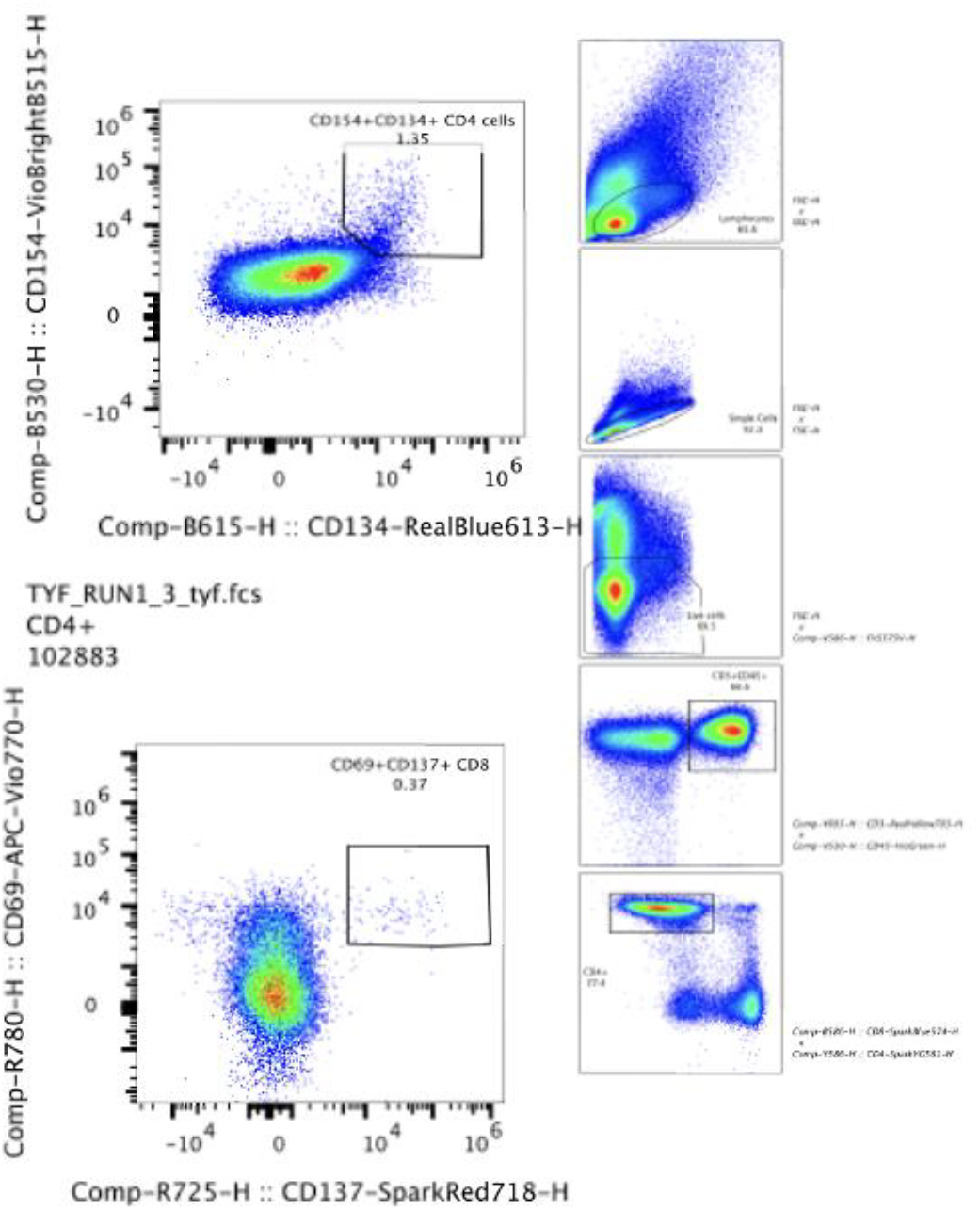
Gating strategy used for identification of phenotypic subpopulations. Representative flow cytometry plots showing the sequential gating steps used to define activated (AIM^+^) T cells and their phenotypic subsets. AIM^+^ CD^4+^ T cells were identified based on co-expression of CD134 and CD154. Gating was applied consistently across all samples and stimulation conditions using FlowJo. Plots shown are representative of the YFV-stimulated condition.

**Supplementary figure 2.**
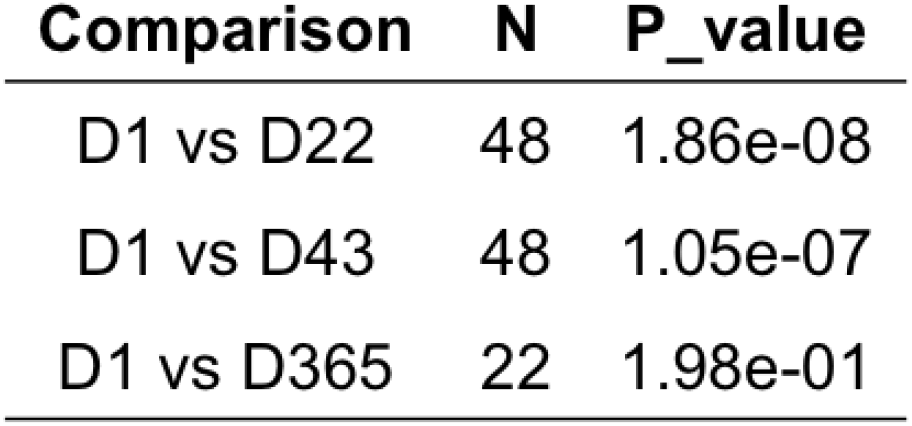
Statistical comparison of Δ AIM% in CD^4+^ T cells between timepoints following yellow fever peptide stimulation. P-values were calculated using paired Wilcoxon signed-rank tests comparing each post-vaccination timepoint (D22, D43, D365) to baseline (D1) within matched donors (N = 48 for D22 and D43; N = 22 for D365).

**Supplementary figure 3.**
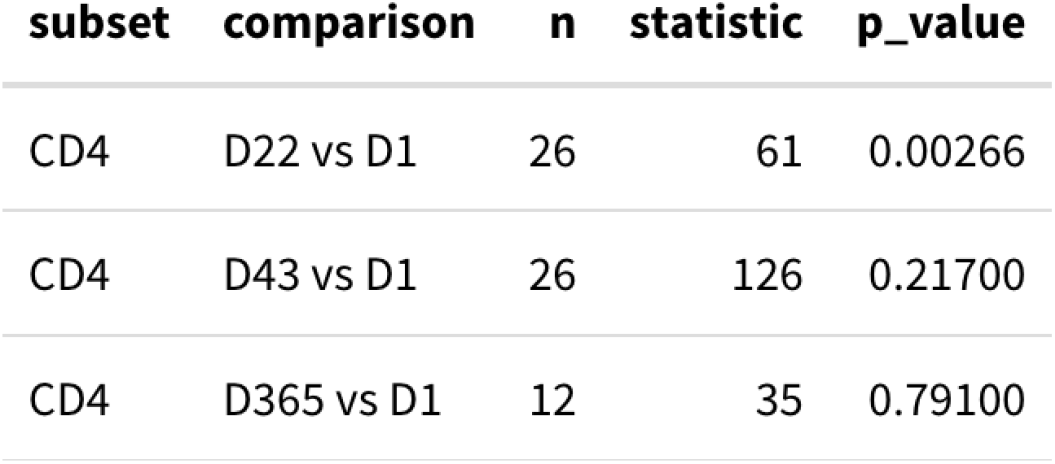
Wilcoxon signed-rank tests comparing Shannon entropy at post-vaccination timepoints versus baseline (D1) for AIM^+^ CD^4+^ repertoire. Pairwise comparisons were performed to evaluate changes in Shannon entropy from D1 to D22, D43, and D365 within the CD^4+^ AIM^+^ subset. The table presents test statistics and unadjusted p-values for each within-subset comparison.

**Supplementary figure 4.**
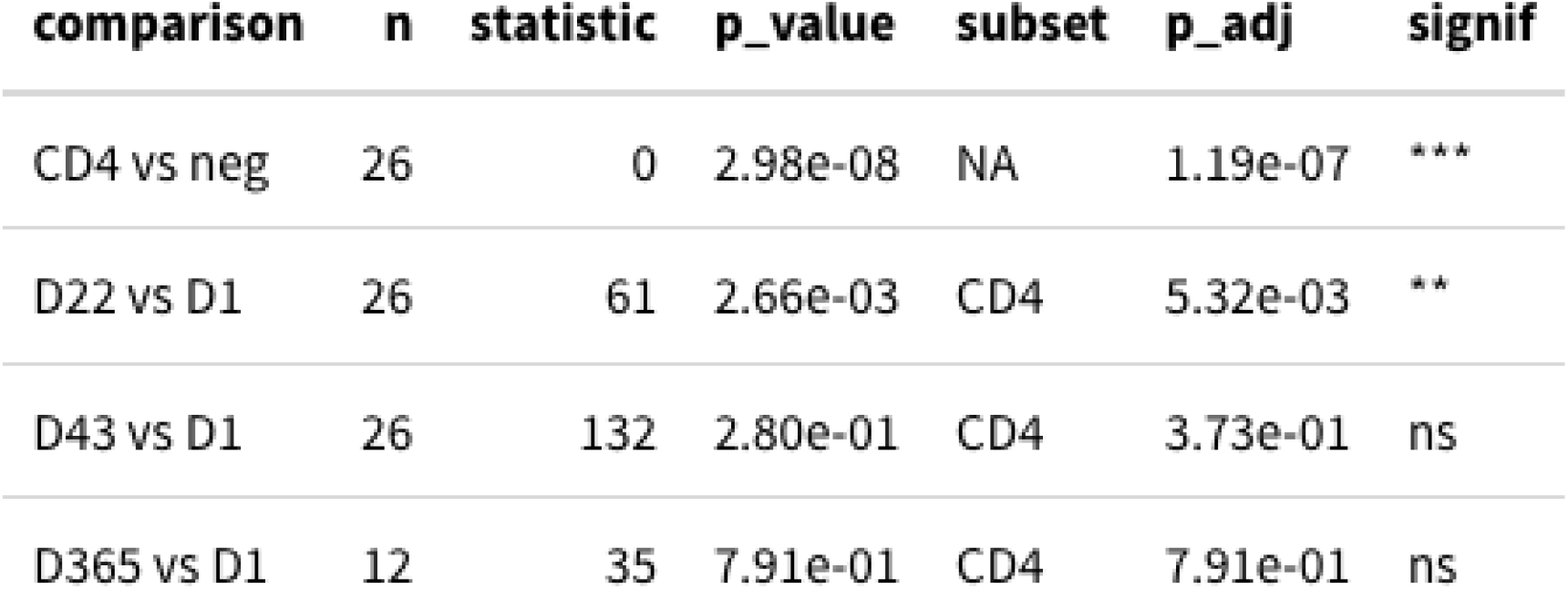
Benjamini-Hochberg corrected p-values for Shannon entropy in AIM^+^ CD4 T cells across timepoints and versus background. P-values from Wilcoxon signed-rank tests were corrected for multiple testing using the Benjamini-Hochberg method. This table summarizes adjusted p-values for (i) CD^4+^ AIM^+^ versus AIM^−^ background, and (ii) entropy changes across timepoints compared to baseline. Asterisks denote statistically significant differences after correction (adjusted p < 0.05).

**Supplementary Figure 5.**
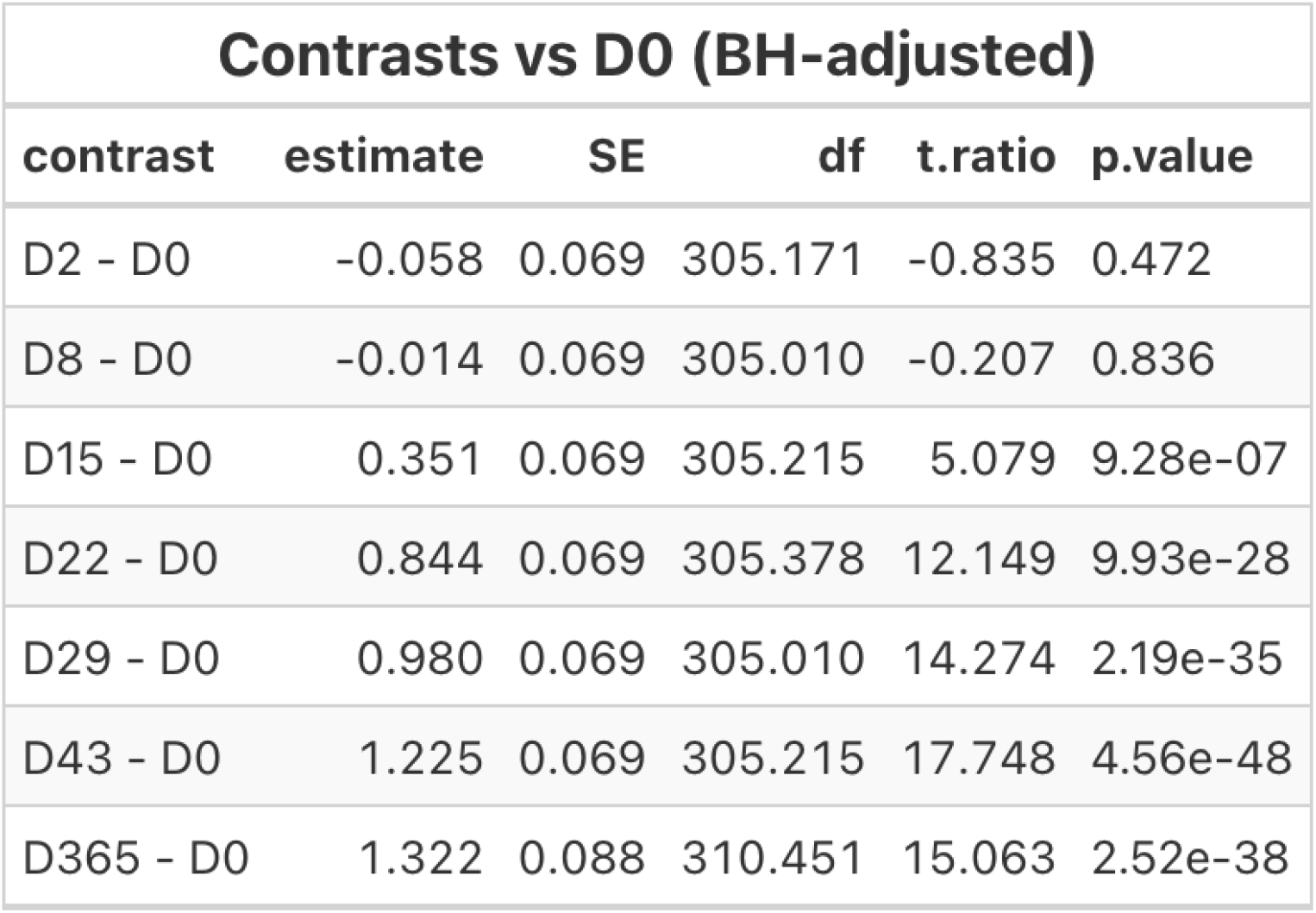
Benjamini–Hochberg corrected contrasts of YFV-specific IgG levels versus baseline (D0). Linear mixed-effects models (LMM) were fitted to log10(FI−Bkgd^+^1) YFV IgG values with participant as a random effect and timepoint as a fixed effect. Estimated marginal means were compared to baseline (D0) using treatment-versus-control contrasts. Resulting p-values were corrected for multiple testing using the Benjamini–Hochberg method. The table reports contrast estimates, standard errors (SE), degrees of freedom (df), t-ratios, and adjusted p-values.

**Supplementary Figure 6.**
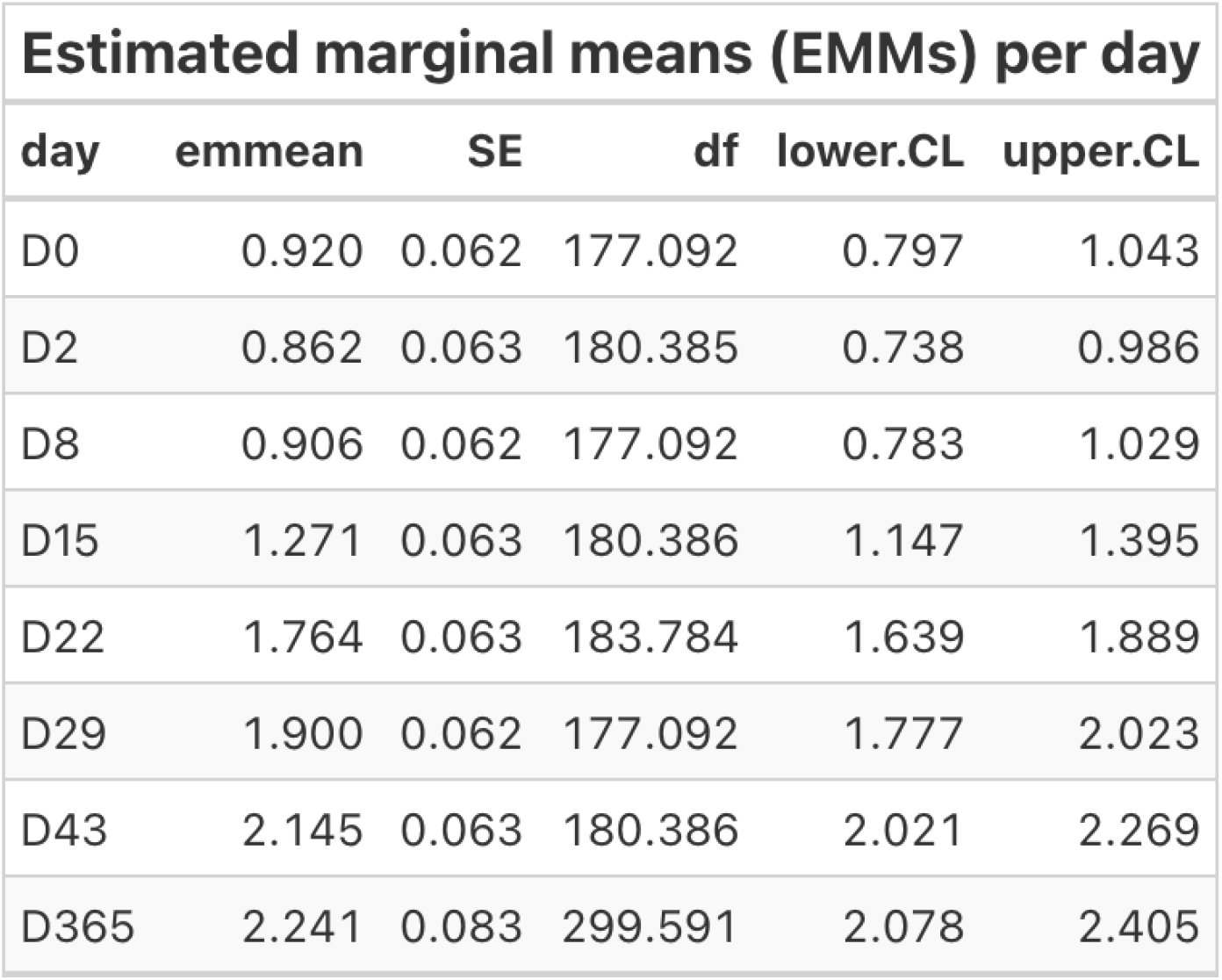
Estimated marginal means (EMMs) of YFV-specific IgG levels across timepoints. Estimated marginal means were derived from the full linear mixed-effects model fitted to log10(FI−Bkgd^+^1) YFV IgG values, accounting for repeated measurements within participants. The table presents model-based mean estimates per timepoint, associated standard errors (SE), degrees of freedom (df), and 95% confidence intervals (lower.CL, upper.CL).

**Supplementary figure 7.**
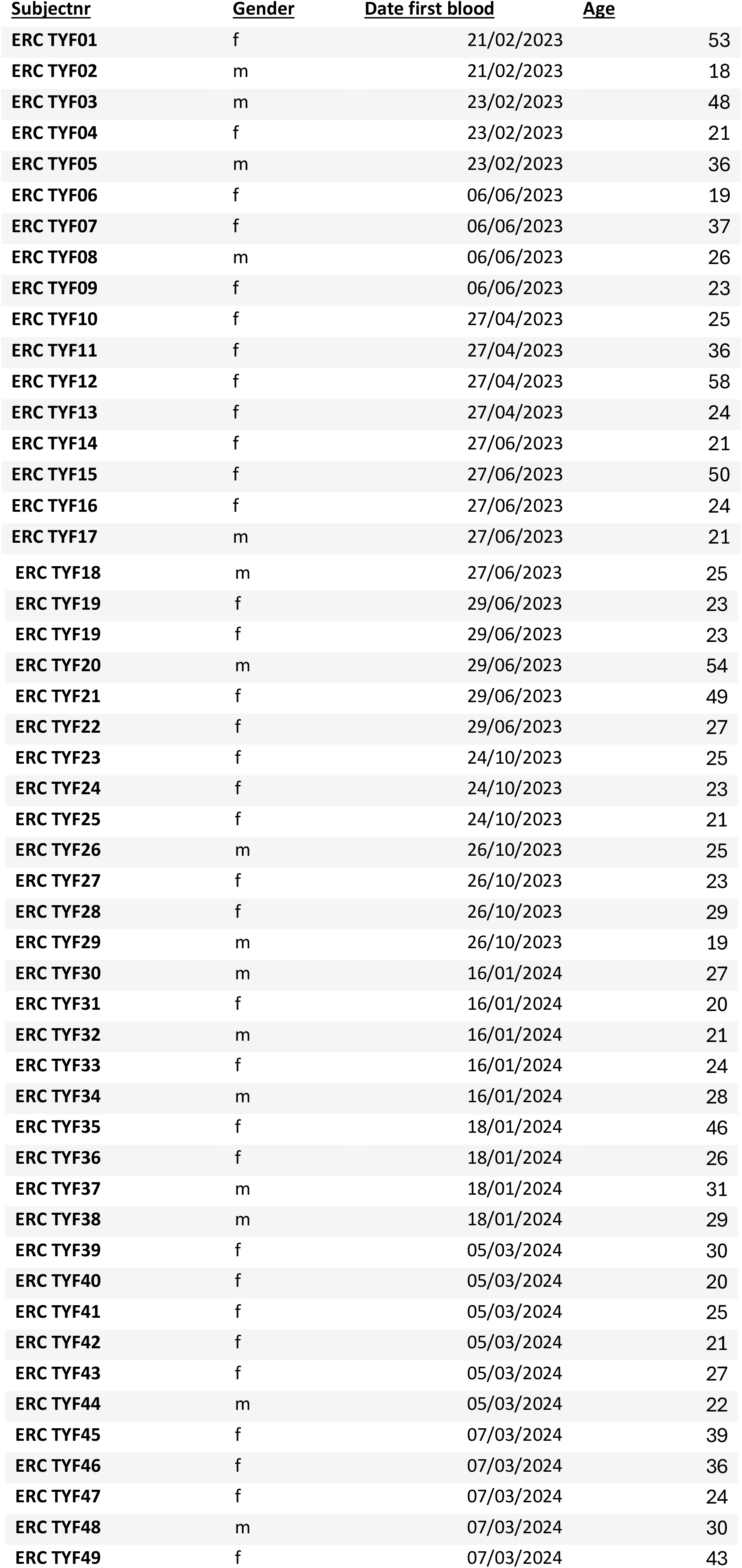
Demographic information for yellow fever vaccine study participants. The table lists subject identifier codes, gender, date of first blood draw, and age. All data were pseudonymized in accordance with ethical and data protection standards.

**Supplementary figure 8.**
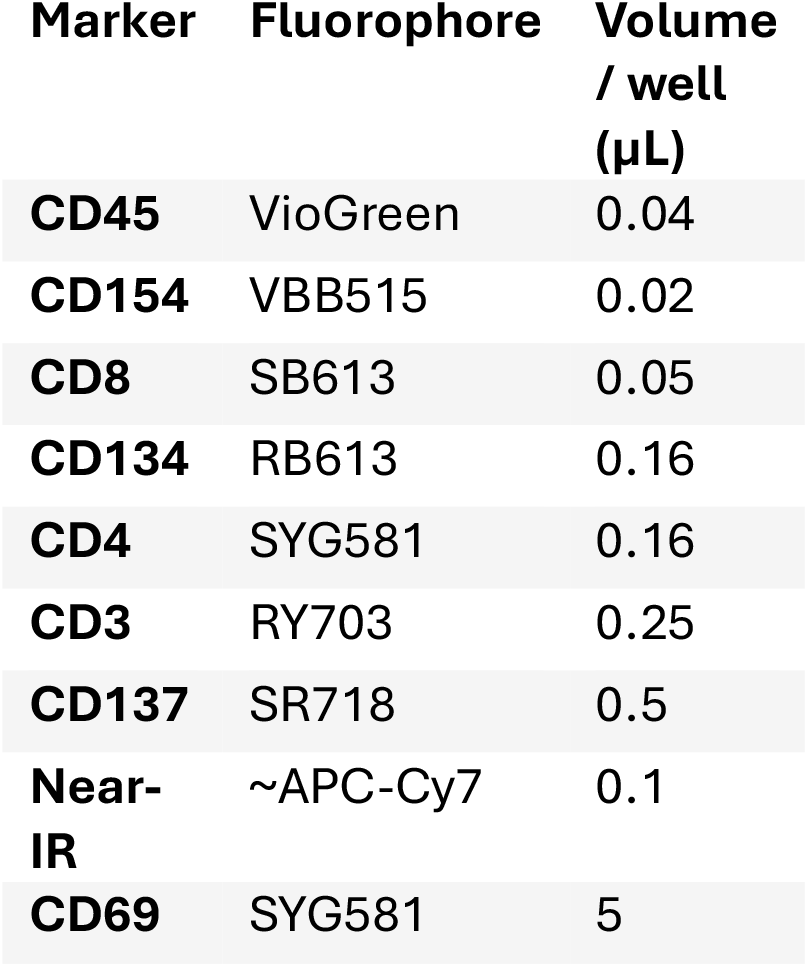
Markers used for cell sorting with Cytek® Aurora CS

**Supplementary figure 9.**
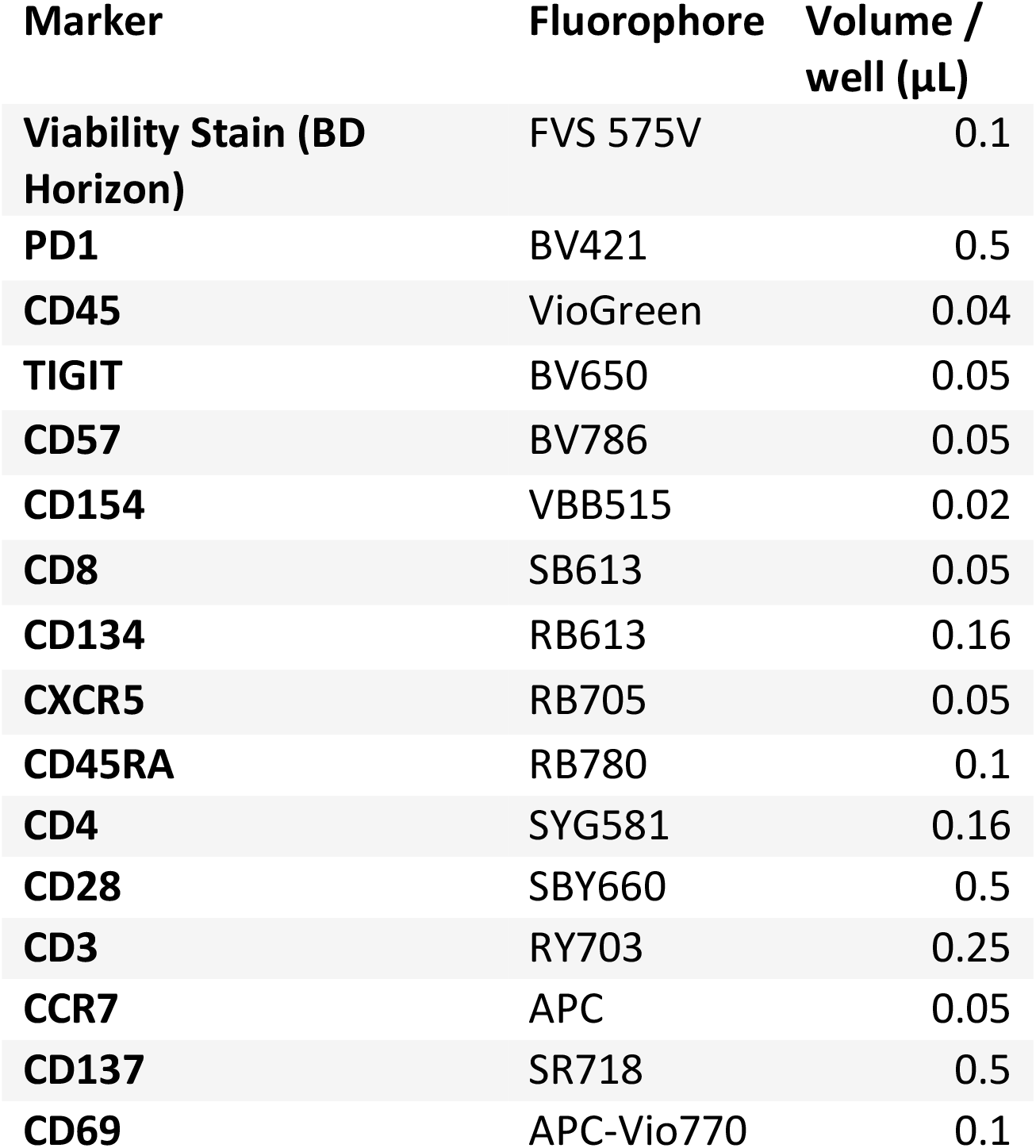
Markers used for flow cytometry analysis with the Quanteon Novocyte Flow Cytometer

